# Protenix-v2: Broadening the Reach of Structure Prediction and Biomolecular Design

**DOI:** 10.64898/2026.04.10.717613

**Authors:** Yuxuan Zhang, Chengyue Gong, Jinyuan Sun, Jiaqi Guan, Milong Ren, Song Xue, Hanyu Zhang, Wenzhi Ma, Zhenyu Liu, Xinshi Chen, Wenzhi Xiao

## Abstract

Advances in biomolecular modeling have broadened the range of problems addressable by structure prediction and design models. Here, we present results from **Protenix-v2**, a system spanning high-accuracy structure prediction and biomolecular design. On the structure prediction side, Protenix-v2 achieves antibody-antigen success rates with up to 13-point gains over Protenix-v1, while **5-seed performance surpasses previous 1000-seed results**. On the design side, Protenix-v2 demonstrates a **100% target-level success rate** in novelty-controlled VHH-Fc campaigns, reaching hit rates up to 48%. Crucially, the model enables hit discovery on **difficult GPCR targets** with hit rates of 16%–88% (VHH-Fc) and up to 50% (mAb) under 16–30 testing budgets per target. Resulting hits show high **developability** and **diversity**. Beyond antibody tasks, we report improved ligand-related plausibility and successful cross-variant SARS-CoV-2 spike RBD mini-binder design. These results establish Protenix-v2 as a robust and powerful model for accelerated drug discovery.

## 1 Introduction

Characterizing biomolecular interactions is central to both mechanistic understanding and molecular discovery. In practice, this requires models that can resolve not only folded structures but also interfaces and the quality of candidate interactions across diverse molecular settings. Recent all-atom systems have substantially broadened what can be modeled in this way, extending high-accuracy prediction from proteins to more general biomolecular complexes relevant to drug discovery [1, 8, 11, 14, 19–21, 28, 33, 34].

In parallel, generative design systems have advanced from general protein generation toward more controlled design, including epitope-conditioned and antibody-format binders [2, 4, 6, 12, 13, 17, 23, 25, 26, 29–31, 39]. These developments make zero-shot hit discovery increasingly plausible, with recent systems beginning to show encouraging breadth across target novelty, antibody format, developability, and target classes [5, 27, 29].

Here, we present Protenix-v2, a biomolecular modeling system spanning structure prediction and molecular design. On the structure side, Protenix-v2 shows clear gains on antibody-antigen structure prediction, together with an additional update in ligand-related plausibility. On the design side, this report moves beyond the protein-binder emphasis in prior PXDesign work [32] toward zero-shot antibody design, developability assessment, and hit discovery on challenging targets such as G protein-coupled receptors (GPCRs), a therapeutically central but experimentally difficult target class for antibody discovery because the accessible extracellular epitopes are often small and conformationally flexible [5, 27]. The key results are:

- **Significant Gains in Antibody-Antigen Structure Prediction**: Protenix-v2 improves across three antibody-focused benchmark collections, showing up to 13-point gains over Protenix-v1 at DockQ *>* 0.23 and comparably large gains in the stricter DockQ *>* 0.8 regime. Crucially, the model demonstrates a massive leap in sampling efficiency: its 5-seed performance notably surpasses previous 1000-seed results.
- **High-Success Zero-Shot Antibody Design**: On novelty-controlled targets, Protenix-v2 achieves a 100% target-level success rate in the current panel, with biolayer interferometry (BLI)-confirmed hit rates ranging from 2% to 48%. The resulting hits show exceptional developability and diversity, with pass rates of 100%, 98%, and 93% in thermostability, self-interaction, and polyreactivity assays, respectively, while spanning multiple structural clusters.
- **Effective Hit Discovery on Challenging Targets**: On difficult GPCRs with small and flexible exposed epitopes, Protenix-v2 yields BLI-confirmed hits in both VHH-Fc and mAb formats. Despite a highly limited testing budget of only 16–30 designs per target, the model achieves hit rates of 16%–88% in VHH-Fc and up to 50% in mAb.

We also report two additional updates: (1) improved ligand-related plausibility on recent protein-ligand benchmarks and (2) dual-specific binders against both prototype and Omicron RBDs with nanomolar-scale *K*_D_. These results establish Protenix-v2 as a robust and powerful model for accelerated drug discovery.

## 2 Methods

Protenix-v2 operates in two primary modes: (1) to predict and rank biomolecular structures, and (2) to generate and prioritize candidate binders for experimental validation. To ensure rigorous evaluation and prevent data leakage, Protenix-v2 is trained without wwPDB [3] entries released on or after 2021-09-30. This training cutoff aligns with established conventions in recent models [8, 28, 34], ensuring that all reported results in structure prediction and zero-shot design are assessed against truly novel or held-out data.

Relative to earlier Protenix iterations [8, 33], Protenix-v2 incorporates architectural refinements and training optimizations while retaining the same input-output setting. For molecular design, Protenix-v2 enables flexible, target-conditioned generation that supports both precise epitope-targeting and site-agnostic design. Its capabilities span diverse protein-binder classes, ranging from miniproteins and modular antibody formats such as variable domain of heavy chain of heavy-chain antibody (VHH), and variable fragment (Fv) including variable domain of heavy chain (VH) and compatible variable domain of light chain (VL). In antibody settings, Protenix-v2 empowers users with granular control over binding regions: complementarity-determining region (CDR) loops can be independently assigned with specified length ranges, while predefined frameworks or scaffolds can be integrated into the design specification to guide the generation process.

## 3 Antibody-Antigen Structure Prediction

Antibody-antigen complexes represent a stringent test for structure-based discovery. While their modeling is therapeutically vital, predicting their interface geometry remains exceptionally challenging [18, 35]. Figure 2 summarizes the performance of Protenix-v2 across three specialized antibody benchmark collections: PXMeter-AB [16], FoldBench-AB [37], and AF3-AB [15]. The evaluation includes AlphaFold3 [1] when published results are available [34, 37], Boltz-1 [36] as a representative baseline for recent all-atom models, and OpenFold3-preview2 (OF3p2) [34] as a very strong open-source comparator. Compared to these baselines and the earlier Protenix-v1, Protenix-v2 demonstrates a substantial improvement trend. At the DockQ *>* 0.23 threshold, Protenix-v2 achieves absolute success rate gains of 9 to 13 percentage points over Protenix-v1 across three collections. Notably, these improvements extend into the high-precision regime: gains at DockQ *>* 0.8 are comparably large, representing a significant relative leap over earlier models. Such advancements bring a wider array of antibody-antigen targets into a practically useful quality range for downstream drug discovery.

The performance advantage is more pronounced when considering inference-time scaling. While Figure 2 summarizes the 5-seed results, Figure 3 illustrates success rate curves up to 1000 seeds. Antibody-antigen prediction was previously shown to benefit from larger inference budgets [1, 8, 34]; however, Protenix-v2 consistently outperforms competitors across the entire explored seed range. Remarkably, Protenix-v2 at only 5 seeds already exceeds the performance of Protenix-v1 at 1000 seeds, indicating a clear gain in efficiency.

In direct comparisons, Protenix-v2 moves from approximate parity with AlphaFold3 to a clear leading position in antibody-antigen modeling.

## 4 Zero-shot Antibody Design

The design results reported here move beyond the protein-binder focus of prior PXDesign work [32] to achieve robust zero-shot antibody generation. To evaluate Protenix-v2 in settings that stress both target novelty and structural complexity, we established a comprehensive evaluation panel spanning soluble targets and challenging membrane proteins to simulate real-world discovery workflows.

- *Soluble Targets*: Rather than reproducing the full target panels from prior studies [23, 29], we first restricted attention to antigens that were available for immediate experimental follow-up, and then sampled from that in-stock pool to cover several published novelty regimes and target classes. The resulting panel spans:
  - Low-Homology Monomers (BoltzGen targets [23]): From the available pool, we included monomeric targets satisfying the very-low-homology setting emphasized by BoltzGen, namely proteins that are highly dissimilar to any earlier protein in the PDB under a 30% sequence identity threshold.
  - Novelty-Filtered Targets (Chai-2 targets [29]): We also included targets from the available pool after excluding antigens and homologs in SAbDab released prior to our training cutoff, following the novelty-filtering logic used in Chai-2.
  - Dimeric Target Case Study: We included VEGF-A to evaluate the model’s design capability on non-monomeric complexes and to probe performance beyond the monomer-only regime.

- *Challenging Membrane Proteins (GPCRs)*: We included multiple GPCR targets, a therapeutically important receptor class whose accessible extracellular epitopes are often small and conformationally flexible, making them difficult settings for both de novo design and traditional antibody discovery. The current hit rates, developability, and diversity results are summarized in Figures 1 and 4 to 6. Shared experimental details for these assays are summarized in Section B.

**Figure 1.**
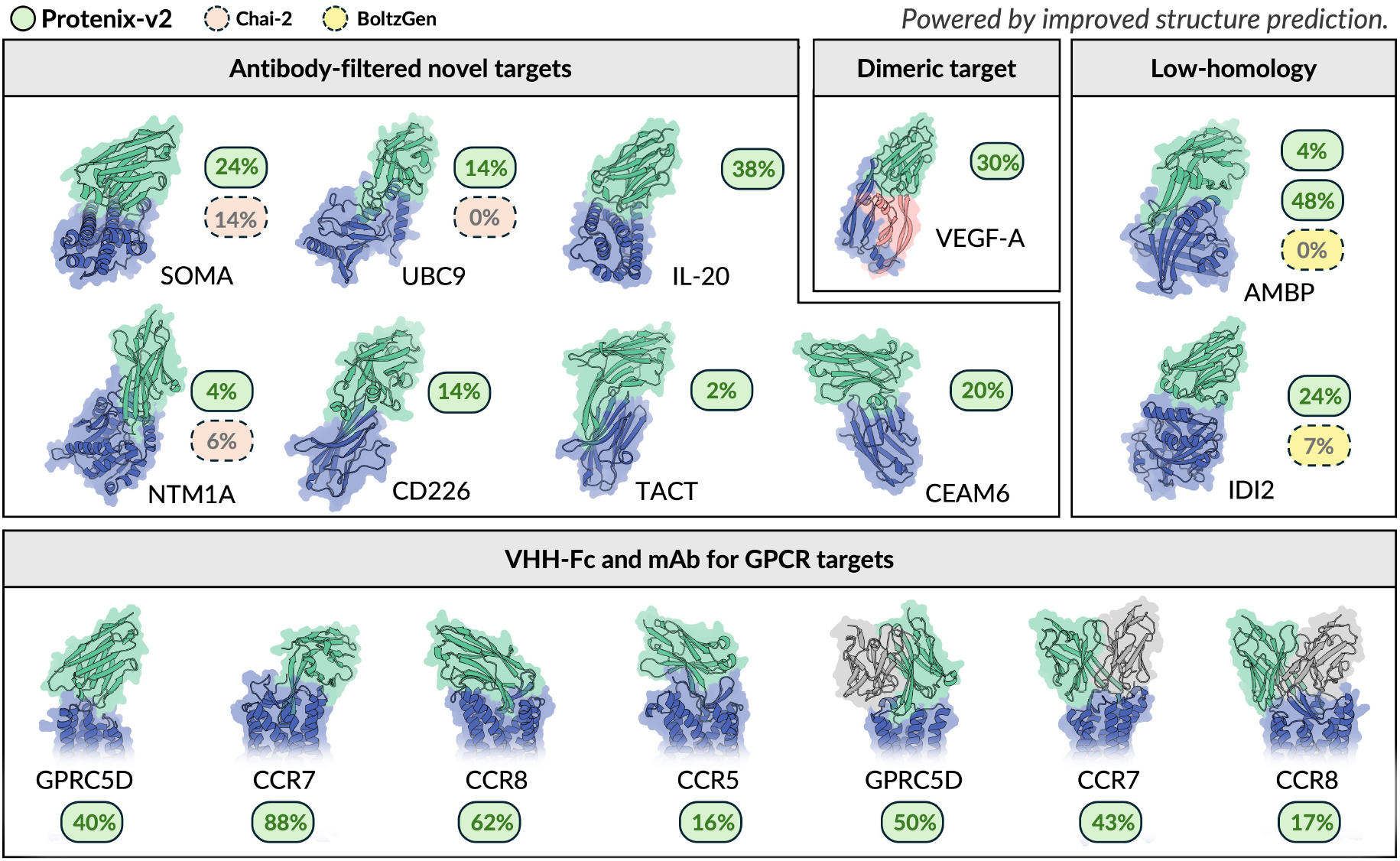
Zero-shot antibody design hit rates, building on the structure-prediction gains (Figure 2).

**Figure 2.**
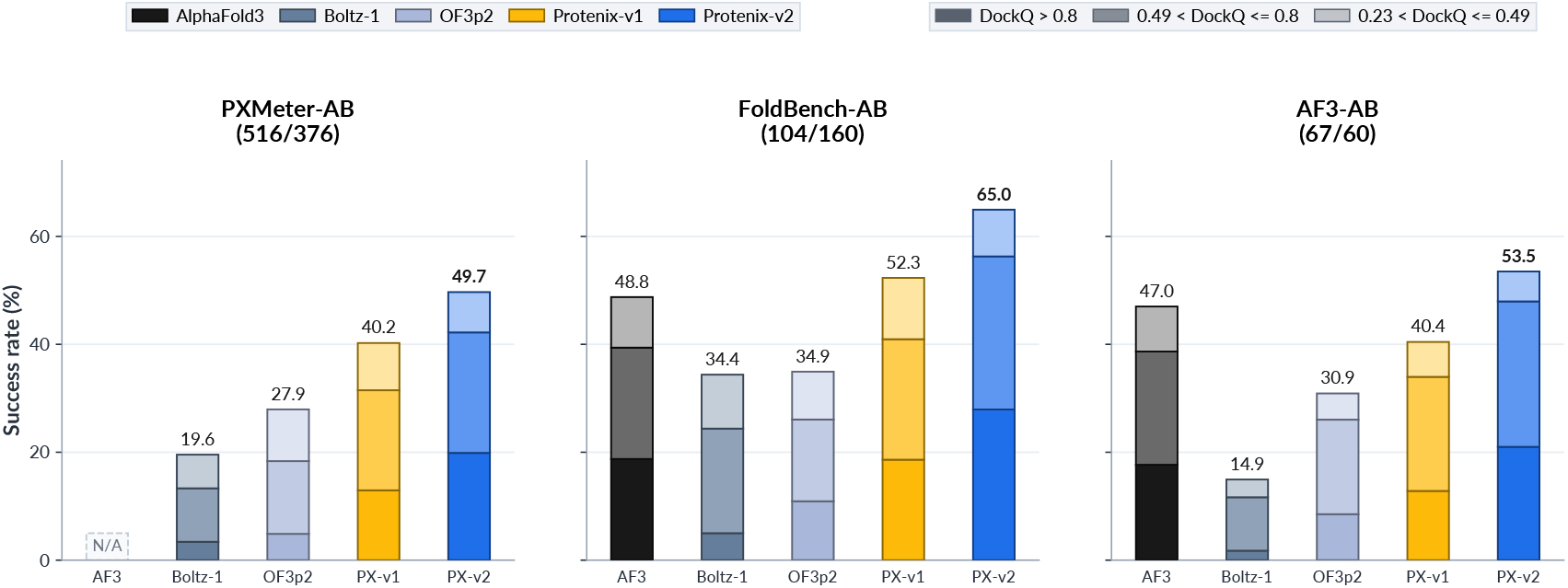
Benchmarking Protenix-v2 (PX-v2) on Antibody-Antigen Interface Prediction. Success rates are reported across three benchmark collections using the ranked top-1 prediction at 5 seeds. In each panel title, the parenthesized pair is written as (number of entries/number of clusters). Bars are stacked cumulatively: the total height represents the success rate at DockQ *>* 0.23, while darker layers denote subsets achieving DockQ *>* 0.49 (medium quality) and DockQ *>* 0.8 (high quality). Boltz-1 and AF3 FoldBench-AB results are from Xu et al. [37] and we apply ranking score selector to the rest models for fair comparison. AF3-AB results are from The OpenFold3 Team [34].

**Figure 3.**
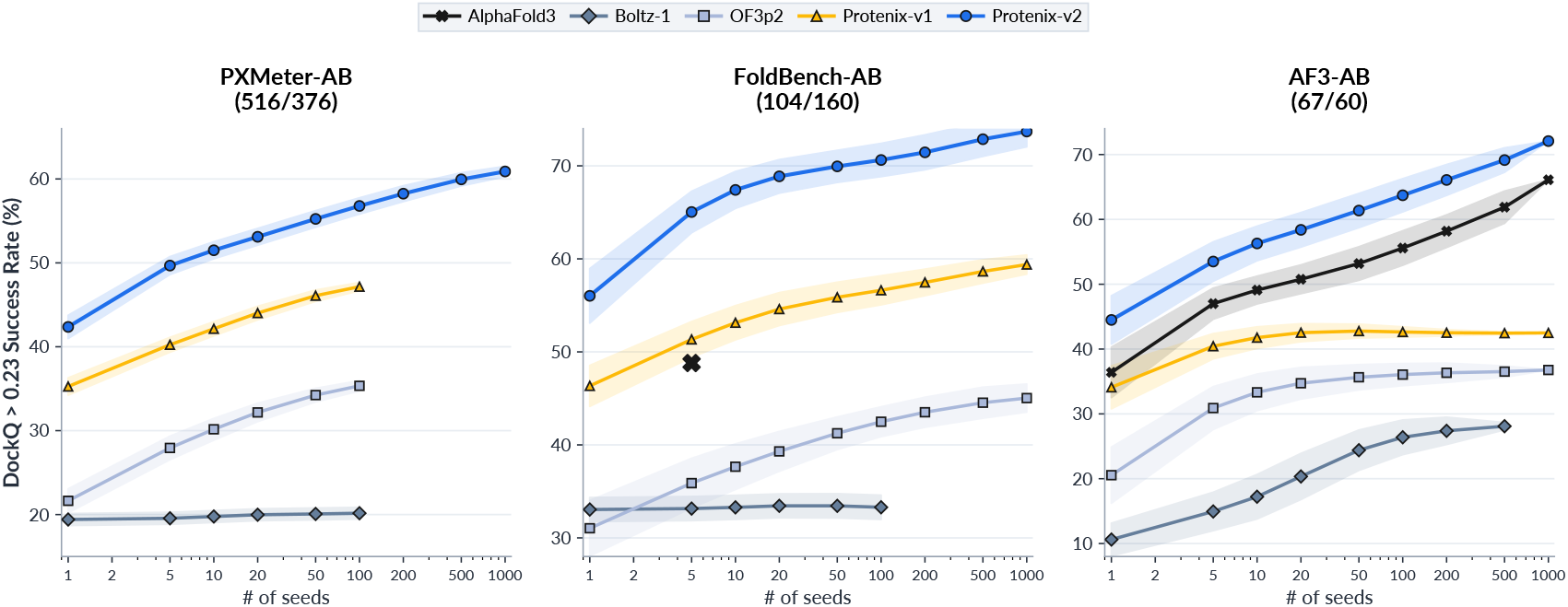
Superior Sampling Efficiency and Inference-Time Scaling. DockQ success rates are plotted as a function of the number of seeds. Shaded bands indicate bootstrap standard deviation. In FoldBench-AB, AlphaFold3 is available only at 5 seeds and is therefore shown as a single marker.

**Figure 4.**
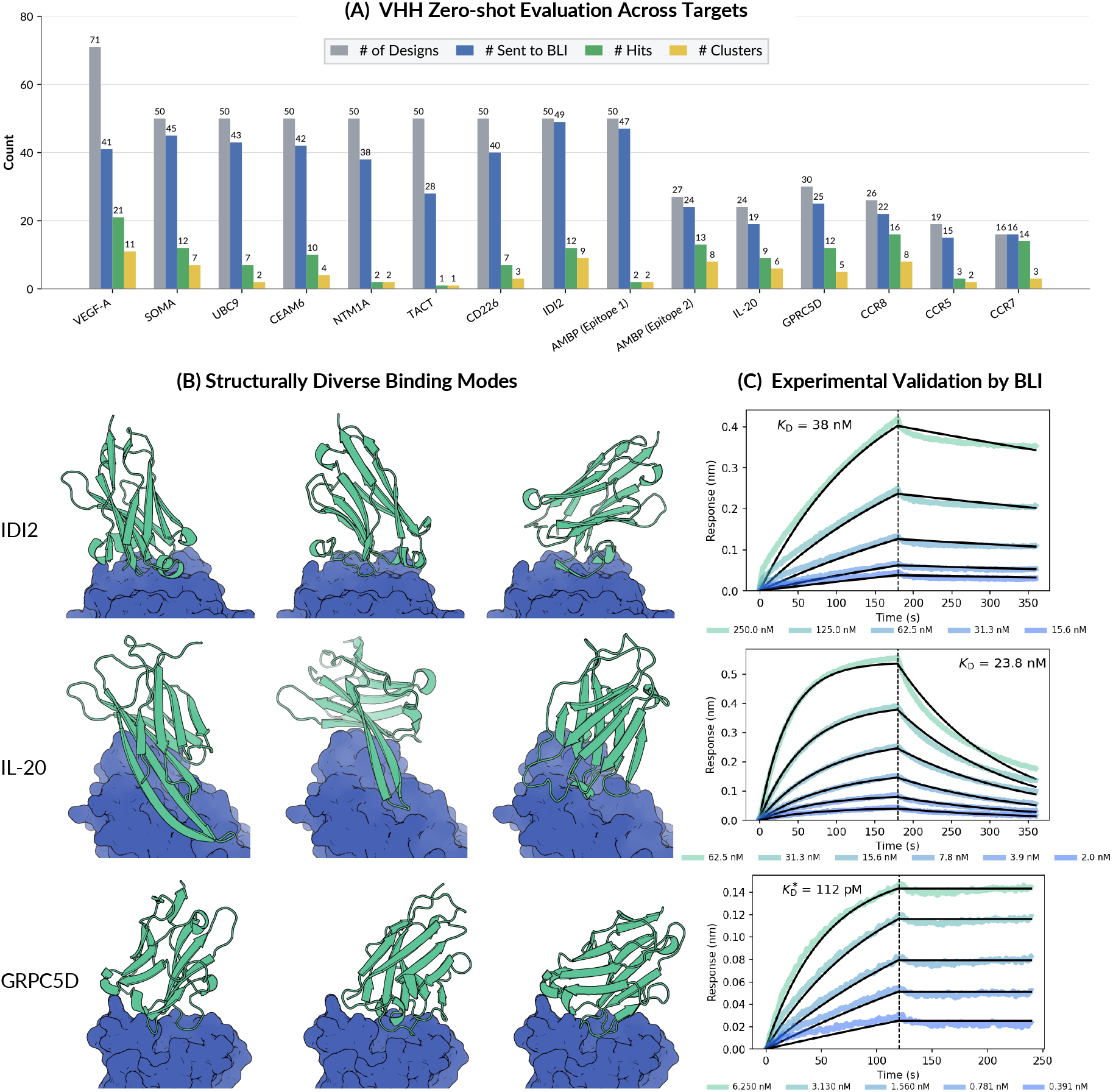
VHH zero-shot evaluation, structural diversity, and kinetic characterization. (A) Statistics across the VHH panel, including the number of raw designs, the subset sent to BLI, the number of confirmed hits, and the number of structural clusters among the hits. (B) Representative binding poses from distinct structural clusters for selected successful campaigns. (C) Examples of BLI sensorgrams for VHH binders. The 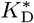 of GRPC5D is measured under avidity conditions due to the antigen’s native dimeric state.

**Figure 5.**
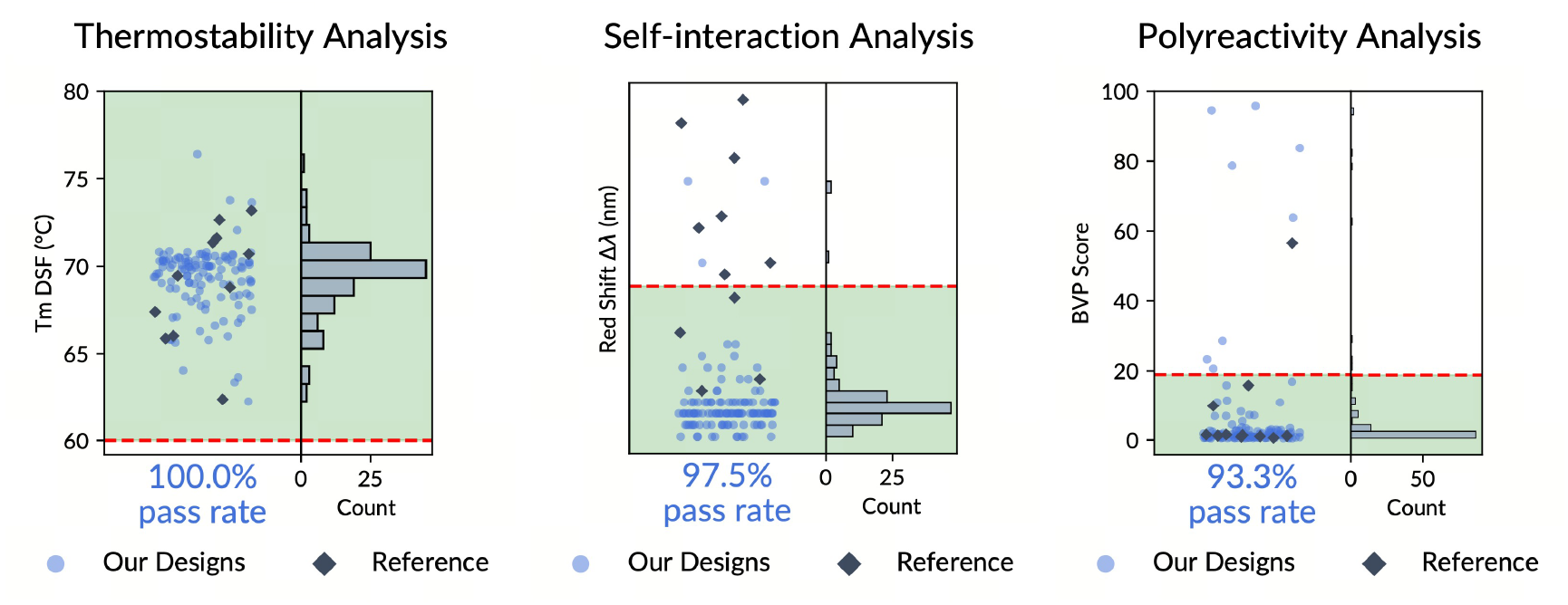
Developability-related experimental readouts. The three panels summarize thermostability, self-interaction, and polyreactivity, with pass rates of 100.0%, 97.5%, and 93.3%, respectively.

**Figure 6.**
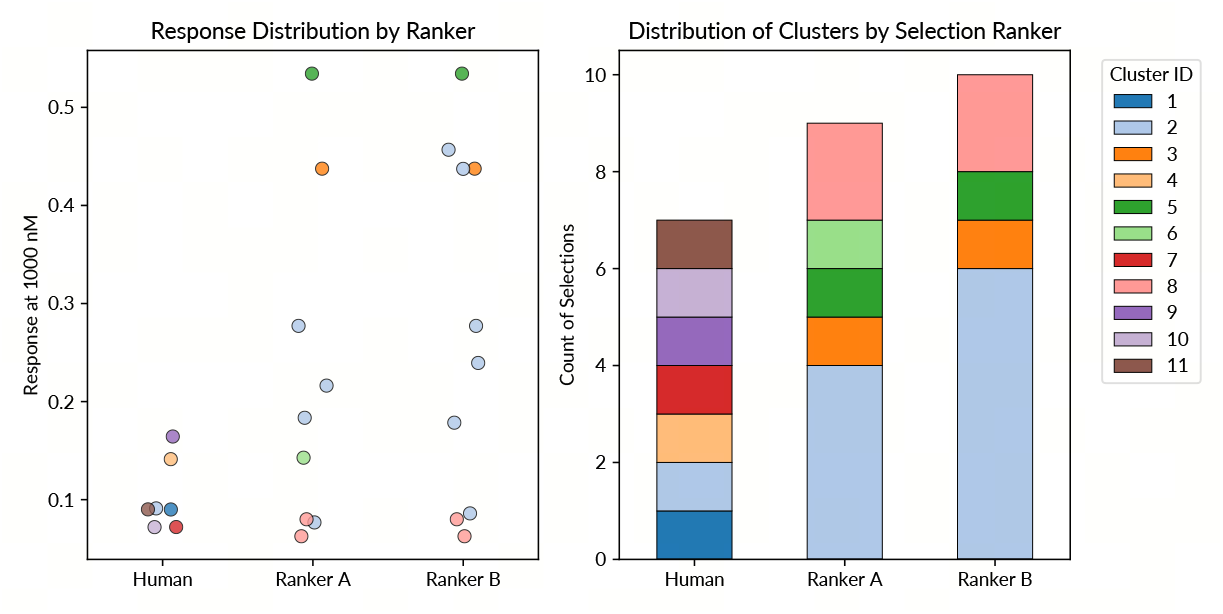
VEGF-A ranking comparison between human and model-based selection. Left: response at 1000 nM for experimentally confirmed binders selected by a human expert and by two model rankers integrated in Protenix-v2.

### Precision and Efficiency on Soluble Targets

The top panels of Figure 1 and Figure 4 (A) summarize the results of VHH designs targeting soluble proteins. Across this diverse evaluation, Protenix-v2 achieves a 100% target-level success rate, meaning at least one experimentally confirmed binder was discovered for every antigen tested. For novelty-filtered soluble targets, BLI-confirmed VHH-Fc hit rates range from 2% to 48%, attaining higher hit rates on most overlapping targets. Direct comparisons are shown only where published VHH baselines are available, since some overlapping Chai-2 targets were reported only in scFv format.

### Epitope choice affects design difficulty

This is clearly illustrated by our results on AMBP. When targeting two different epitopes on the same AMBP protein, Protenix-v2 yielded hit rates of 4% and 48%, respectively. This variance highlights that the choice of surface patch can alter the difficulty of de novo design. Consequently, we determined that aligning our epitope selection for novelty-filtered targets with the methodology used in the Chai-2 study [29], specifically targeting native ligand-binding interfaces.

### Effective Hit Discovery on GPCR Targets

GPCRs are a particularly demanding test for antibody design: they represent a high-value therapeutic target class, yet the exposed receptor surface is often limited, flexible, and poorly suited to conventional antibody discovery workflows [5, 27]. In this setting, we evaluated Protenix-v2 in a zero-shot regime with 16–30 tested designs per target and in two distinct antibody formats: VHH-Fc and full-length mAb (monoclonal antibody). Despite this limited experimental budget, Protenix-v2 produces appreciable hit rates in both formats. As summarized in the bottom panel of Figure 1 and Figure 4 (A), across four GPCR targets, the VHH-Fc campaigns reached hit rates of 16%, 62%, 40%, and 88%, while the corresponding mAb campaigns reached 0%, 17%, 50%, and 44%. These results indicate that the model can remain sample-efficient even on difficult membrane-protein targets, while also transferring effectively across formats. For GPRC5D, the lowest *K*_D_ of VHH-Fc designed by Protenix-v2 was 112 pM (at the bottom of Figure 4 (C), measured under avidity conditions due to the antigen’s native dimeric state).

### Developability and Diversity

Binding alone is not sufficient for practical antibody discovery. Successful antibody hits must also show favorable developability, including drug-like biophysical properties. We therefore examined whether the resulting molecules satisfy standard developability criteria, while also achieving meaningful affinity and structural diversity. Protenix-v2 designs exhibit strong pass rates in these measured developability assays: 100% for thermostability, 98% for self-interaction, and 93% for polyreactivity (Figure 5). Furthermore, the collection of successful binders spans multiple structural clusters when grouped by antigen-aligned framework RMSD at 4 Å (Figure 4 (A)), indicating that the recovered solutions are not confined to a single repeated pose family. Representative examples of these diverse binding modes are shown in panel (B) of Figure 4. Together, these measurements suggest that the zero-shot hits are not only experimentally recoverable binders, but also promising discovery starting points from the perspectives of biophysical quality, binding-mode diversity, and biochemical potency.

### Ranking Capability Beyond Human Intuition

VEGF-A provides a useful ranking-focused case study because it is dimeric and therefore falls outside the monomer-only settings emphasized in prior benchmarks. We compared a human expert with two rankers integrated in Protenix-v2 (Ranker A and Ranker B) on approximately 300 designed VHH candidates, each selecting 30 for experimental validation. Ranker A and Ranker B identified 9 and 10 binders, respectively, with 5 overlapping hits. Each model also contributed unique binders (4 for Ranker A and 5 for Ranker B), indicating complementary strengths. Also, the human expert identified 7 binders, all of which were unique and showed no overlap with either model. These results suggest that the rankers capture binding-relevant features that are largely orthogonal to human intuition. As shown in Figure 6, model rankers selected hits with higher response at 1000 nM compared to human, but human selected more diverse hits than models.

The model rankers recover more high-response binders, while the human expert tends to select weaker-response candidates. Right: structural-cluster composition of the selected binders. Human selections span a broader set of clusters, whereas the model rankers concentrate more strongly on a smaller number of productive clusters.

## 5 Expanded Capabilities: Chemical Fidelity and Multi-Variant Design

Beyond antibody-specific tasks, Protenix-v2 also delivers advancements in small-molecule modeling and multi-target design, addressing critical needs in chemical realism and therapeutic breadth.

### 5.1 Enhancing Ligand Plausibility via Training-Free Guidance

Ensuring the physical plausibility of predicted small-molecule structures is critical in ligand binding pose prediction, because strong agreement with the reference under conventional geometric metrics does not necessarily guarantee a chemically realistic local geometry. Errors such as chirality inversion or incorrect bond lengths can directly affect the credibility of predicted interactions, the interpretation of ligand strain, and the usefulness of the model output in expert review.

Structural plausibility is commonly assessed using the PoseBusters validity criterion [7]. Under this criterion, some models achieve strong pass rates by introducing inference-time steering [20, 36]. However, our manual case review shows that good performance on the standard validity checks does not imply that predicted ligands satisfy other basic forms of chemical plausibility that are also important to medicinal chemists. For example, we observe failures in properties such as twisted amide groups, even when the corresponding predictions pass the original validity filter, as shown in figure 8. Therefore, in the new version of PXMeter (v1.1.0), we extend the standard PoseBusters validity criterion with additional checks on planarity around sp2 carbon centers, planarity of amide groups, and non-planarity at sp3 carbon and nitrogen centers, enabling a more comprehensive assessment of ligand pose.

Building upon our base model, we develop the Protenix-v1-TFG and Protenix-v2-TFG variants, drawing inspirations from Boltz-1x [36], training-free guidance [38], projected diffusion [9], and classical constraint-enforcement schemes such as the SHAKE algorithm [22] in molecular dynamics. In particular, these variants impose constraints on chirality, planarity, torsional geometry, and pairwise distances, thereby guiding the generative process toward ligand conformations that better satisfy fundamental stereochemical and geometric requirements.

We evaluate our methods and several comparison baselines on the PXM-22to25-Ligand test set, which comprises protein–ligand complexes released between 2022 and 2025 and therefore provides a challenging test of model performance on recent structures. Performance is evaluated using a joint success metric requiring both a pocket-aligned ligand RMSD below 2 Å and a validity pass. We report results under both the original PoseBusters criterion and a revised criterion that additionally includes the metrics described above. As expected, Table 1 shows that imposing the stricter validity criterion lowers absolute success rates for almost all models. Even under this stricter evaluation, Protenix-v2-TFG reaches 60.46%, markedly improving over Boltz-1x (53.96%) and approaching the 62.86% achieved by Boltz-2x. Protenix-v1-TFG also attains a strong 61.11%, indicating that the proposed TFG variants remain highly effective. Notably, while Protenix-v2-TFG is already comparable to Boltz-2x, the latter is shown for reference only, because its 2023 training cutoff means that part of this evaluation set may overlap with its training data.

**Table 1.** Ligand-related plausibility on PXM-22to25-Ligand (623 test entries, 250 clusters), reported as the joint success rate requiring both pocket-aligned ligand RMSD *<* 2 Å and validity to pass. “PoseBusters” uses the original validity definition, while “Revised” applies the stricter definition described in the main text.

| Model | RMSD & Validity SR (%) |  |
| --- | --- | --- |
|  | PoseBusters | Revised |
| Boltz-2<sup>†</sup> (train overlap) | 51.31 | 51.25 |
| Boltz-2x<sup>†</sup> (train overlap) | 63.51 | 62.86 |
| Boltz-1 | 35.56 | 34.38 |
| Boltz-1x | 55.59 | 53.96 |
| Protenix-v1 | 48.37 | 48.14 |
| Protenix-v1-TFG | 61.11 | 61.11 |
| Protenix-v2 | 50.42 | 49.22 |
| Protenix-v2-TFG | 60.53 | 60.46 |

To understand which validity checks are most informative under the revised evaluation, we next examine the individual PoseBusters-style checks. Figure 7 shows that the largest gaps occur in planarity around sp2 centers and non-planarity at sp3 centers, with the TFG variants consistently remaining closer to ground truth than the Boltz-1x and Boltz-2x baselines. The near-100% pass rate of ground-truth structures on these checks further justifies the inclusion of the newly introduced metrics. These results further show that existing models still exhibit significant weaknesses in generating chemically plausible small-molecule geometries, even when they perform well under the original criterion.

**Figure 7.**
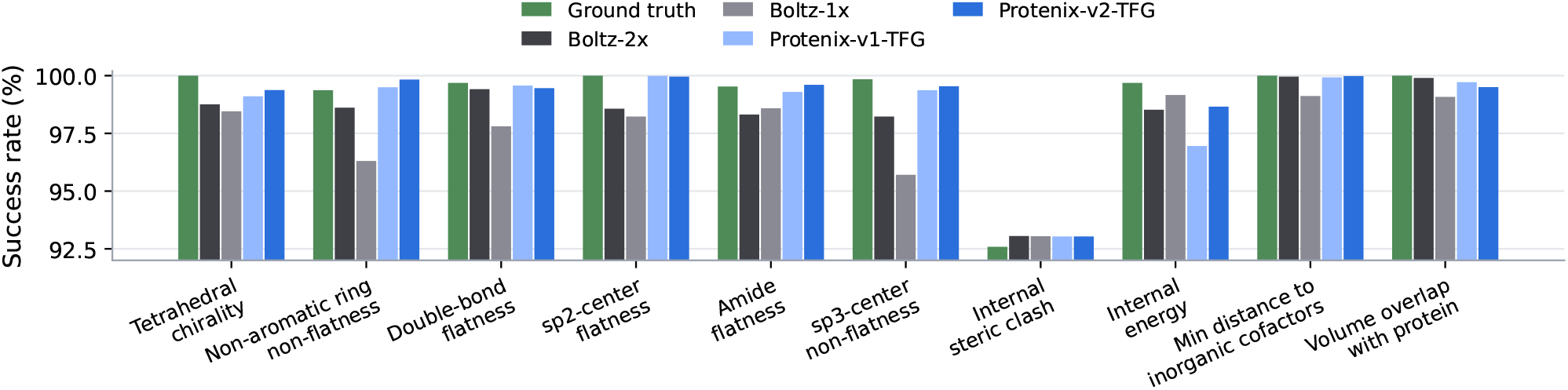
Selected breakdown of revised-validity success rates across ten informative PoseBusters-style subitems for ligand-containing structures.

**Figure 8.**
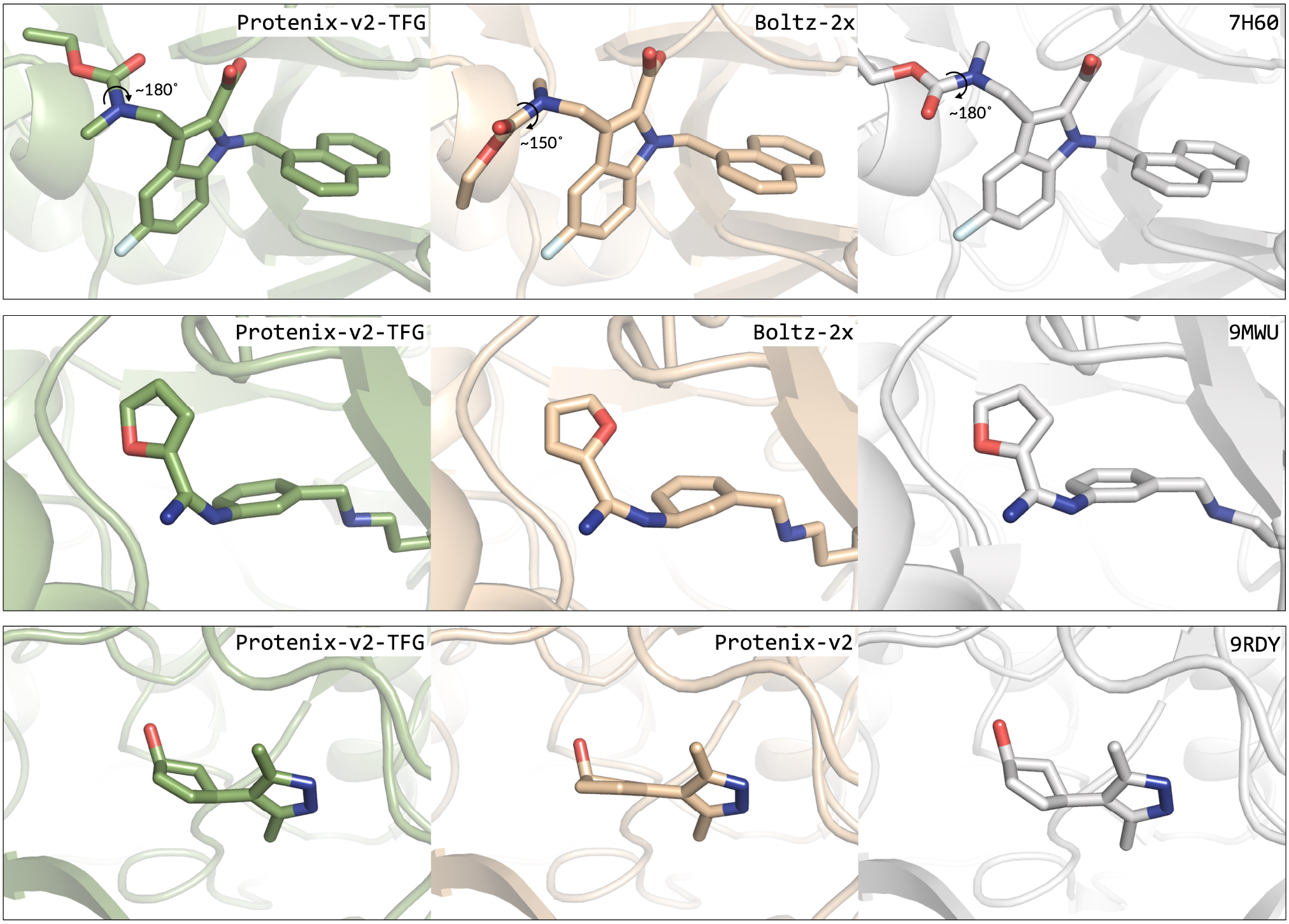
Case studies of protein–ligand structure prediction demonstrating the effectiveness of the revised physical plausibility evaluation criterion with the TFG variants passing the check successfully while Protenix-v2 or other baseline model fail. Top: Boltz-2x predicts a twisted amide group. Middle: Boltz-2x predicts a distorted aromatic ring that passes the original check but is detected by the sp2 flatness check. Bottom: Protenix-v2 predicts an incorrectly planar geometry around an sp3 carbon center.

### 5.2 Dual Binding Across Prototype and Omicron SC2RBD Variants

Breadth-oriented binder design provides a stringent test for Protenix-v2, as effective candidates must retain binding across variants with substantial sequence and epitope differences.

Previous PXDesign mini-binders showed weak-to-none binding to the RBD of SARS-Cov-2 Omicron B.1.1.529. In contrast, when RBDs of both prototype and Omicron were jointly provided as inputs, Protenix-v2 generated 2 dual-binding mini-binders out of 4 tested designs. Both exhibited nanomolar-scale *K*_D_ against the RBDs of the prototype and Omicron B.1.1.529 variants. Based on the predicted structures, we performed preliminary analysis to understand the design as shown in the Figure 9. At the structural level, the broad-spectrum binder exhibits potential compensatory mechanisms. In the prototype complex, residue R13 of the binder may form polar interactions with the main-chain carbonyl of L492 or the side-chain amide of Q493 on the RBD. Simultaneously, the side-chain amide of N102 on the binder may also form a polar interaction with the side-chain amide of Q493. In the predicted complex structure of the binder with the Omicron variant, the Q493R mutation on the RBD introduces steric hindrance and charge alterations. The main-chain carbonyls of L103 and E101 on the binder likely participate in polar interactions with R493. Additionally, the side chain of N102 on the binder reorients to establish potential specific interactions with the main-chain carbonyl of S494 on the RBD.

**Figure 9.**
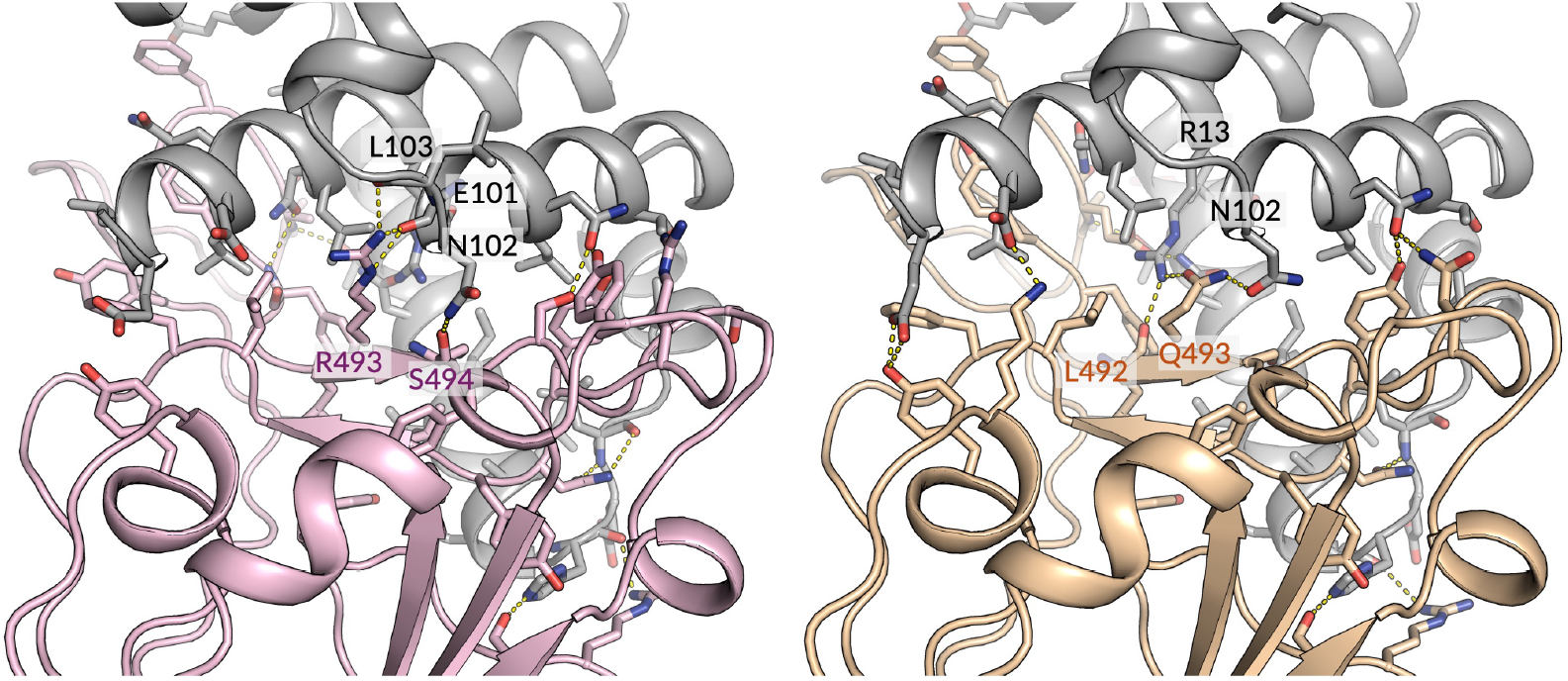
Detailed view of the interaction interfaces. Left panel is the binder complex with Omicron RBD, the right panel is the binder complex with prototype RBD. Binder is colored in grey. In both panels, key residues at the interface are represented as stick models and labeled with matching background colors. Yellow dashed lines indicate polar contacts between the chains.

**Figure 10.**
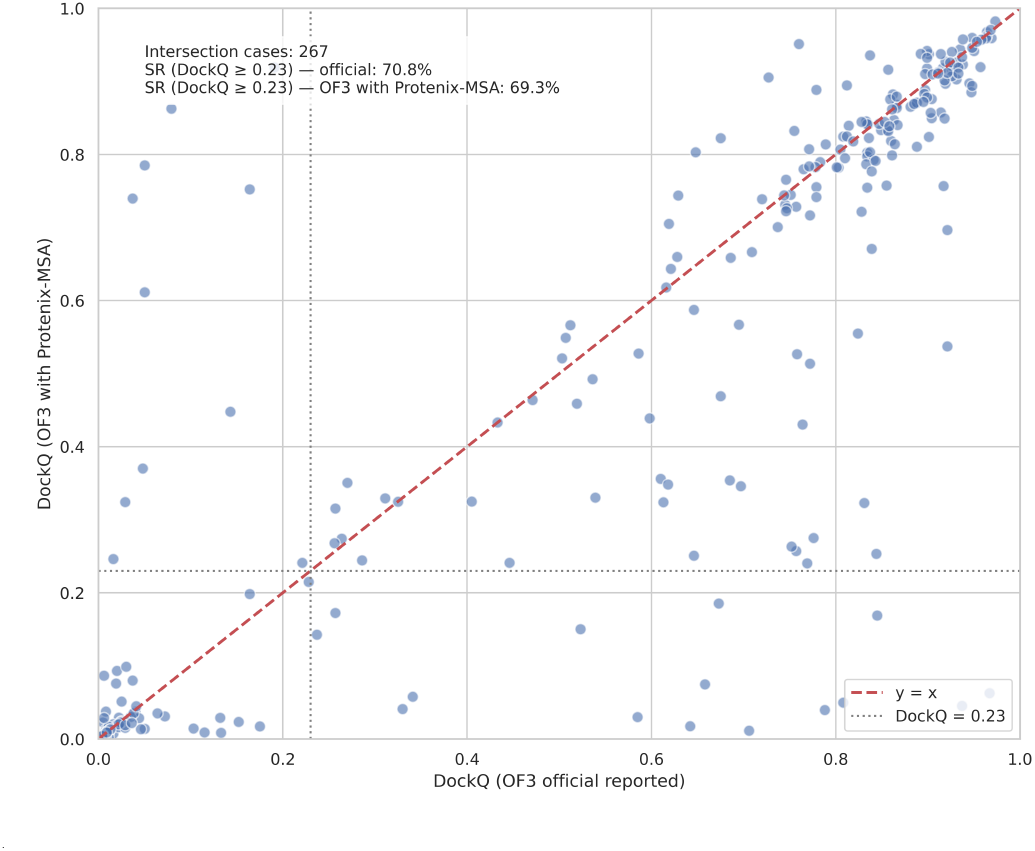
Impact of MSA/Template Choices on OpenFold3 DockQ Success Rate. We compare DockQ success rate (SR; DockQ ≥ 0.23) on FoldBench protein–protein interfaces between the public OpenFold3-preview2 FoldBench result and our OF3p2 inference using an MMseqs-derived MSA pipeline without templates. On a common set of 267 interfaces, the resulting DockQ SR is 69.29% in our setting versus 70.79% in the public FoldBench result, i.e. about 1.5 points lower. This indicates a measurable but modest effect from the MSA/template setting difference.

These results highlight the ability of Protenix-v2 to design binders with cross-variant activity, which is critical for developing therapeutics resilient to viral evolution. Moreover, cross-reactivity across related targets is also highly desirable in drug development, as it facilitates evaluation in animal models and supports more efficient preclinical translation.

Table 2 summarizes the current dual-binding SC2RBD results. Experimental details for this breadth-oriented assay are summarized in Section B.

**Table 2.** Affinity of Protenix-v2-designed SC2RBD mini-binders to both prototype and Omicron B.1.1.529 RBDs. Lower *K*_D_ indicates stronger binding.

| Design | Prototype RBD $K_D$ (nM) | Omicron B.1.1.529 RBD $K_D$ (nM) |
| --- | --- | --- |
| SC2RBD Binder 4 | 10.1 | Weak binding, $K_D$ not measured |
| SC2RBD Binder 14 | 10.3 | No binding |
| Design 1 | 250 | 148 |
| Design 2 | **201** | **146** |

## 6 Summary

Protenix-v2 strengthens both structure prediction and biomolecular design in settings that matter directly for therapeutic discovery. On the structure side, the clearest gain is in antibody-antigen modeling, where the updated system improves substantially across three antibody-focused benchmark collections and preserves favorable scaling as the inference budget increases. The model also improves ligand-related plausibility on a recent protein-ligand benchmark, with gains in both pose accuracy and validity.

On the design side, Protenix-v2 supports zero-shot VHH discovery on novelty-controlled targets, with BLI-confirmed hit rates ranging from 2% to 48%, together with encouraging developability pass rates and diverse binding poses. The same capability extends to difficult GPCR targets, where zero-shot campaigns in both VHH-Fc and mAb formats yield high hit rates under limited experimental budgets. We also include a breadth-oriented SC2RBD mini-binder result, where two of four tested designs retain nanomolar-scale binding to both prototype and Omicron B.1.1.529 RBDs. Taken together, these results broaden the practical scope of the Protenix system across antibody-related modeling and design tasks.

## Acknowledgment

This report builds on the cumulative efforts of many contributors across the Protenix series [8, 16, 32, 33]. We first thank Qinru Bai, Zhaolong Li, Yeyu Su, Meijie Deng, Rong Han and Xingang Peng for direct supports to this report, including case studies, internal review, and other support that strengthened the final presentation.

We also thank the contributors of earlier Protenix and PXDesign projects, including Lan Wang, Yanping Yang, Yu Xia, Cong Liu, Yuzhe Wang, Qixu Cai, Chan Lu, Jincai Yang, Ke Zhang, Shenghao Wu, Kuangqi Zhou, Bo Shi, Shaochen Shi, even when they were not directly involved in the current release.

We are similarly grateful to Jiarui Lu, Eric Alcaide, Zehua Chen, Yiran Xu, Jintao Zhu, Bo Qiang, Yiheng Wu, Chaoran Cheng, Fred Peng, Wei Tang, Jiale Zhao, Junxi Mu, Yuyang Zhang, Lu Yan and Yingtian Liu for support across earlier stages of these projects, including interns and other collaborators who assisted with review, analysis, discussion, and related project execution.

Finally, we thank ByteDance for sustained support of this line of work, and we especially acknowledge Hang Li and Liang Xiang for leadership, organizational support, and trust in the Protenix and PXDesign programs.

*April 2026*

*When Beijing meets Seattle*

## Appendix

### A. Benchmark Details

#### A.1 Comparison on FoldBench and PXMeter

This comparison is included as a broader supplemental view of Protenix-v2 beyond the antibody-focused analyses above. FoldBench [37] and PXMeter [16] provide a complementary summary across protein monomers, protein-ligand complexes, and protein-protein interfaces, and are useful for situating the current model against a wider set of recent structure prediction systems.

Table 3 presents a compact side-by-side summary for the two benchmark suites. In this broader view, Protenix-v2 remains competitive across all three interaction categories, while the main-text antibody-focused and ligand-related analyses provide the clearest picture of its strongest practical gains.

**Table 3.** Summary comparison on FoldBench and PXMeter. For each benchmark suite, we report representative top-ranked performance summaries for protein monomers, protein-ligand complexes, and protein-protein interfaces, following the standard metric conventions of the corresponding benchmark. For each benchmark subset, the header reports entries/clusters, i.e. the number of test entries and the corresponding number of clusters. The reported metrics are LDDT, RMSD *<* 2&LDDT *>* 0.8 precentage and DockQ *>* 0.23 success rate for monomer, protein ligand interface and protein protein interface, respectively.

| Model | FoldBench |  |  | PXMeter |  |  |
| --- | --- | --- | --- | --- | --- | --- |
|  | Monomer | Protein-Ligand | Protein-Protein | Monomer | Protein-Ligand | Protein-Protein |
|  | 287/287 | 441/441 | 237/237 | 769/500 | 623/250 | 2126/1806 |
| AlphaFold3 | 88.0 | 62.6 | 71.7 | - | - | - |
| OF3p2 | 86.8 | 56.6 | 66.4 | 85.4 | 47.9 | 63.4 |
| Protenix-v1 | 88.6 | 62.5 | 72.7 | 87.2 | 48.1 | 72.9 |
| Protenix-v2 | 89.1 | 64.0 | 73.0 | 88.0 | 49.2 | 73.9 |

#### A.2 Model Inference Setting

Input files for Protenix, Boltz, and OpenFold3 were generated from mmCIF structures using the gen-input module in PXMeter v1.1.0. Since Boltz and OpenFold3 do not support ligands composed of multiple CCD codes or glycans, these components were excluded from their respective inputs. All models were run with 10 recycling iterations during inference.

For OpenFold3-preview2 (OF3p2), we used an MMseqs2-derived MSA pipeline [24] without templates. This differs slightly from the setting described in the technical report [34], although the public OpenFold3 repository also provides an MMseqs-based MSA pipeline. We checked the practical impact of this difference on FoldBench protein–protein interfaces. On a common set of 267 interfaces, our OF3p2 setup achieves a DockQ success rate of 69.29%, compared with 70.79% in the public FoldBench results, i.e. about 1.5 points lower. We therefore view the mismatch in MSA/template features as real but modest in its effect on overall PPI performance.

### B. Wet-Lab Experimental Details

#### B.1 Recombinant Protein Construction and Design

VHH candidates were formatted as VHH-Fc fusions with a C-terminal human IgG1 Fc (C220S). The designed Fv including VL and VH domains were grafted onto Trastuzumab framework as a full length mAb. To avoid avidity effects due to the natural homodimeric state of VEGF-A, the anti-VEGF-A VHH binders were engineered with a C-terminal His_6_ tag rather than an Fc domain.

#### B.2 Protein Expression and Purification

All recombinant constructs were synthesized and cloned into high-expression mammalian vectors. These proteins were produced using a CHO mammalian cell transient expression system. Cells were cultured in serum-free media in a humidified incubator at 37 °C with 8% CO_2_. Culture supernatants were harvested 5 to 7 days post-transfection and clarified by centrifugation and 0.22 *µ*m filtration.

Fc-tagged proteins, including VHH-Fc and mAbs, were purified using Protein A affinity chromatography with MabSelect resin. Following a PBS wash at pH 7.4, proteins were eluted with 10 mM Glycine at pH 1.5 and immediately neutralized. His-tagged proteins were purified via immobilized metal affinity chromatography using Ni-NTA resin. Purified proteins were subsequently dialyzed or buffer-exchanged into 0.05% PBST or PBS. Protein concentrations were determined by A280 absorbance using a NanoDrop spectrophotometer. The molecular weight and purity under reducing and non-reducing conditions were verified by SDS-PAGE, confirming the integrity of VHH-Fc at approximately 80 kDa, mAb at 150 kDa, and VHH-His at 22 kDa. The monomeric content and homogeneity were further assessed by size-exclusion high-performance liquid chromatography (SEC-HPLC). Only protein batches with a purity of greater than 90% as determined by SEC-HPLC were utilized for subsequent kinetic assays.

For mini-binders, we largely followed the experimental protocol described in our previous report [32]. Strep-tagged constructs were expressed using an *E. coli*-based cell-free system and subsequently purified. Protein concentrations were determined prior to measuring *K*_D_ by multi-concentration BLI.

#### B.3 Binding Kinetics

##### Equipment and Reagents

BLI binding assays were performed using an Octet RH16 instrument (Sartorius) equipped with ProA or ProG Biosensors (Sartorius) to facilitate an antibody-antigen capture format. Histagged RBD proteins were immobilized with NTA sensor. Biotinylated VEGF-A was in immobilized with SA sensor. Assays were prepared in either 96-well or 384-well microplates. Across all experimental stages, 0.05% PBST was utilized as the standard assay and running buffer. Sino Biological catalog number for SARS-CoV-2 B.1.1.529 sublineage BA.2 (Omicron) Spike RBD is 40592-V08H123, and 40592-V08H for prototype SARS-CoV-2 Spike RBD.

##### Antigens

All the antigens were purchased from Sino Biological. The antigen catalog numbers are in Table 4.

**Table 4.** Antigens used in the antibody-design campaigns and the corresponding Sino Biological catalog numbers.

| Antigen | Sino Cat No. |
| --- | --- |
| SOMA | 16122-H07E |
| UBC9 | U224-380H |
| IDI2 | 15292-H07E |
| AMBP | 13141-H08H1 |
| CEAM6 | 10823-H08H-B |
| MTM1A | 11222-H09E |
| TACT | 11202-H08H |
| CD266 | 10565-H08H |
| IL-20 | 13060-HNAE |
| CCR5 | 13022-H91H-NA |
| CCR8 | 11450-H91H-NA |
| GPRC5D | 24447-H73H-NA-B |
| CCR7 | 12112-H95H-NA |
| VEGF-A | 11066-H27H-B |

**Table 5.** Thresholds Defination for Developability Assays.

| Assay | Thresholds | References |
| --- | --- | --- |
| DSF | $T_m > 60\text{ }^{\circ}\text{C}$ | [10] |
| BVP ELISA | Corrected BVP Score $< 5.3$ | [10] |
| AC-SINS | Red Shift $< 11\text{ nm}$ | [10] |

##### Single-Concentration BLI Screening

Initial binding activity of the antibodies was assessed through a single-concentration screening method. All assay steps were performed at an agitation speed of 1000 rpm. For all targets except VEGF-A, the designed antibodies (VHH-Fc or mAb) were immobilized. In detail, ProA or ProG biosensors, depending on the species of control antibodies, were first equilibrated in 0.05% PBST for a 60-second baseline step. The target antibodies (5.0 *µ*g/mL) were subsequently loaded onto the biosensors for 200 seconds to reach a target capture level of approximately 1.8 nm. Following a second 60-second baseline step in 0.05% PBST to remove unbound antibodies, the biosensors were immersed in a 1000 nM antigen solution for 150 seconds to monitor association. Finally, the sensors were moved into 0.05% PBST for 150 seconds to record dissociation. Positive binding candidates were identified based on an established threshold, defined as a binding response greater than the buffer background plus 0.05 nm. Most hits passed this threshold were sent to Multi-concentration *K*_D_ determination, and showed *K*_D_ lower than 1000 nM.

For VEGF-A, the antigen was immobilized on the Streptavidin (SA) Biosensor and the antibodies were used as analytes under the same settings described above.

##### Multi-Concentration *K*_D_ Determination

Antibodies that met the positive screening criteria were advanced to multi-concentration BLI testing to determine precise binding kinetics. The multi-concentration assay maintained a constant agitation speed of 1000 rpm. Following an initial 60-second baseline in 0.05% PBST, antibodies (5.0 *µ*g/mL) were loaded onto the ProA or ProG biosensors for an extended duration of 300 seconds, aiming for a capture signal between 1.94 and 2.0 nm. After establishing a second 60-second baseline in 0.05% PBST, the association phase was measured for 150 seconds by exposing the loaded sensors to a serial dilution of the antigen ranging from 1000 nM to 31.25 nM or lower. The dissociation phase was then tracked for 150 seconds in 0.05% PBST.

For the VEGF-A target, the reversed assay orientation used in the single-concentration screening was maintained, while all other kinetic settings remained the same.

#### B.4 Developability

To comprehensively evaluate the biophysical properties, manufacturability, and potential liabilities of the candidate antibodies, a suite of developability assays was performed. In this study, non-specific binding was assessed using a baculovirus particle (BVP) ELISA, while self-interaction was evaluated using a PEG-optimized affinity-capture self-interaction nanoparticle spectroscopy (AC-SINS) assay. The thermal stability of the candidates was characterized by apparent melting temperature (Tm) via differential scanning fluorimetry (DSF). Eleven reference antibodies from the report of Jain et al. [10], including tabalumab, sarilumab, bavituximab, tremelimumab, guselkumab, dupilumab, teplizumab, brodalumab, motavizumab, inotuzumab, and bimagrumab, were produced and analyzed alongside the designed antibodies to enable calibration of assay passing thresholds across different experimental instruments. All experiments were performed in room temperature unless otherwise specified.

##### Thresholds Defination

For each developability assay, the passing threshold was defined according to the criteria described by Jain et al. [10], with additional calibration based on the reference antibodies included in this study.

##### Polyreactivity

To evaluate the non-specific binding properties of the antibodies BVP ELISA was performed. Briefly, microtiter plates were coated with BVP and incubated overnight at 4 °C. Following a single wash step with wash buffer, the plates were blocked with blocking buffer for 1 hour. After washing three times, primary antibody samples were diluted to a final concentration of 100 nM, added to the wells, and incubated for 1 hour. The plates were then washed six times and incubated with a horseradish peroxidase (HRP)-conjugated Goat Anti-Human IgG-Fc secondary antibody (Sigma, A0170) for 1 hour. After six final washes, Amplex Red substrate (Beyotime, ST010) was added and incubated and protected from light. The fluorescence readout was measured at 565 nm using a SpectraMax i3X multimode microplate reader (Molecular Devices). Besides the reference antibodies above, infliximab, bevacizumab, lenzilumab, and gantenerumab were also used as the standard assay reference. To ensure comparability, the BVP scores were normalized using a linear regression model based on reference antibodies mentioned above from the report by Jain et al. [10].

##### Thermostability

Antibody samples were diluted to a final concentration of 1 mg/mL in PBS (pH 7.0–7.2) and loaded into a FrameStar 384-well skirted PCR plate (Azenta) at a volume of 15 *µ*L per well. Measurements were performed in duplicate using a SuprDSF instrument (Applied Photophysics). A thermal ramp was applied from 20 °C to 100 °C at a heating rate of 1 °C/min, with an integration time of 25 ms. The Tm and the onset of melting (Tm onset) were determined by monitoring changes in intrinsic tryptophan and tyrosine fluorescence at emission wavelengths of 330 nm and 350 nm.

##### Self-Interaction

AC-SINS was utilized to evaluate the self-association of the antibody candidates. First, AffiniPure Goat Anti-Human IgG (Fc*γ* fragment specific) capture antibodies (Jackson ImmunoResearch, 109-005-098) were buffer-exchanged into sodium acetate. The capture antibodies were then mixed and incubated with gold nanoparticles (AuNPs, Zhongkekeyou, ZKKYZW-1-20) at room temperature to allow for conjugation. The AuNP-protein complexes were subsequently blocked by adding poly(ethylene glycol) methyl ether thiol (Sigma, 729140) and incubated at room temperature. Following centrifugation, the complexes were resuspended in PBS. Antibody samples to be tested were diluted to 0.05 mg/mL, mixed with the AuNP-capture antibody conjugates, and incubated at room temperature. The plasmon wavelength shifts of the nanoparticle conjugates were analyzed by detecting the absorbance spectrum using an EPOCH2 microplate reader (Agilent).

